# Dynamic regulation of CD45 by tetraspanin CD53

**DOI:** 10.1101/854323

**Authors:** V. E. Dunlock, A. B. Arp, E. Jansen, S. Charrin, S. J. van Deventer, M. D. Wright, L. Querol-Cano, E. Rubinstein, A. B. van Spriel

**Affiliations:** Department of Tumor Immunology, Radboud Institute for Molecular Life Sciences, Radboud University Medical Center, 6525 GA Nijmegen, Netherlands; Sorbonne Université, INSERM, CNRS, Centre d’Immunologie et des Maladies Infectieuses, CIMI-Paris, Paris, France; Department of Immunology, Monash University, Alfred Medical Research and Education Precinct, Melbourne, Victoria 3004, Australia

## Abstract

T cells are central to the adaptive immune response, playing a role in both the direct and indirect killing of pathogens and transformed cells. The activation of T cells is the result of a complex signaling cascade, initiated at the T cell receptor (TCR), and ending with the induction of proliferation. CD45, a member of the protein tyrosine phosphatase family, is one of the most abundant membrane proteins on T cells and functions by regulating activation directly downstream of the TCR. As a result of alternative splicing, CD45 can be expressed in multiple isoforms, naive T cells express the CD45RA isoform, while activated T cells gain expression of CD45RO, which has been proposed to increase signaling. Though the importance of CD45 in TCR signaling, proliferation and cytokine production is well established, little is known about the regulation of CD45 activity. We discovered that the immune-specific tetraspanin CD53 directly affects the stability and function of CD45RO in T cells.

We have identified CD53 as a T cell co-stimulatory molecule in primary human and murine cells. Furthermore, we have shown that the absence of CD53 leads to an altered CD45 isoform expression as a result of decreased CD45RO stability on the cell surface. This instability was accompanied by increased mobility as measured by FRAP.

Together, this indicates that CD53 functions as a stabilizer of CD45RO, and therefore as a positive regulator of TCR signaling at the T cell surface. Our data provides novel insight into the role of tetraspanins in the regulation of immune signaling and may provide a new avenue for the regulation of T cell signaling.

## Introduction

The adaptive immune response relies heavily on the activation, proliferation and differentiation of T cells, in order to function properly. This activation is the direct result of T cell receptor (TCR) binding to peptide-MHC (pMHC) complexes on the surface of antigen presenting cells (APC). The interaction between these two surface receptors initiates the formation of the immunological synapse, a dynamic interface through which APCs and T cells exchange information through sustained signaling. Downstream of the TCR, signaling is propagated through different protein tyrosine kinases (PTKs), including Lck and ZAP70. The ensuing signaling pattern is the result of a tightly controlled balance between protein phosphorylation and dephosphorylation, both of which, depending on the context, can activate or inhibit signaling molecules. Protein tyrosine phosphatases (PTPs) are very important in the TCR signaling process as a counterpart to the PTKs, but still understudied. This vital role is reflected in their expression pattern, with more than half of the known PTPs reported to be expressed by T cells_1_. One of the most important PTPs expressed by T cells is the transmembrane protein CD45, which occupies approximately 10 % of the T cell surface and can be found on all nucleated hematopoietic cells_2, 3_. In the absence of CD45, murine T cell development and activation are severely impaired, highlighting the importance of this protein for normal T cell function_4_. Thymocyte maturation is blocked at the immature CD4+ CD8+ double-positive stage in CD45-deficient mice_4_. Furthermore, mutation of CD45 in humans has been linked to severe combined immunodeficiency (SCID) and autoimmune disorders_5, 6_.

CD45 is a large single pass transmembrane glycoprotein (180-220 kDa) responsible for dephosphorylating the tyrosine kinase Lck at position Y505 in T cells, thereby relieving the auto-inhibition of Lck_7_. Activated Lck is then able to directly phosphorylate the TCR and ZAP70, and thereby initiates the T cell signaling cascade. CD45 is encoded by 35 exons, producing multiple isoforms through selective inclusion of exons 4-6, which are involved in alternative mRNA splicing_8_. At the protein level, exons 4, 5 and 6 are referred to as A, B and C respectively, and the resulting protein products are given names with letters specifying the included exons with the letter R preceding, to indicating that the protein is restricted to the exons denoted. Thus, the largest isoform, containing all three exons, is known as CD45RABC while the smallest isoform which lacks all three exons is denoted as CD45RO. Five main isoforms are reported to be expressed on human T cells, namely, CD45RO, CD45RB, CD45RAB, CD45RBC and CD45RABC. In mice 4 major isoforms have been identified, these include CD45RO, CD45RB, CD45RBC and CD45RABC_8, 9_.

Naïve human T cells express mainly the high molecular weight (MW) isoforms containing the A exon (CD45RA+ cells). This expression pattern is lost upon antigen encounter and activation and activated T cells gain expression of the low MW isoform CD45RO (CD45RO+ cells). Murine T cells have been reported to undergo a similar switch upon activation, by losing an isoform expressing exon B, and gaining expression of a lower molecular mass isoform lacking exon B, likely CD45RO based on the reported molecular weight_10_. Despite the importance of CD45 in the immune system, the regulation of CD45, especially isoform-specific regulation, remains poorly understood. Here we show for the first time that the RO isoform of CD45 in T cells is regulated by interaction with the tetraspanin protein CD53.

Tetraspanins are a family of 4-transmembrane proteins involved in the organization of proteins on the cells surface. They interact with each other and with partner proteins, forming membrane structures known as tetraspanin enriched microdomains (TEMs). Tetraspanins have been shown to play important roles in numerous cell functions including, adhesion and migration, fusion and cell signaling_11, 12, 13, 14_. Though immune cells express many tetraspanin proteins, there are only two known immune-specific tetraspanins: CD37 and CD53. We have previously shown that CD53 plays a role in the regulation of B cell signaling, but CD53 function in T cells remains unknown_14_.

Here we report a novel molecular interaction between CD45RO and CD53 in both human and murine T cells. Our findings indicate that tetraspanin CD53 is required for stabilization of CD45RO on the T cell surface, and CD53-deficiency impairs the activation and proliferative capacity of T cells.

## Results

### CD53 is a novel partner of CD45

In order to investigate the role of CD53 in T cell function, we performed immunoprecipitations (IPs) followed by mass spectrometry to identify novel interacting partners. CD53 and CD81 were immunoprecipitated from Raji B cell lysates under varying conditions (Figure 1A). As shown, a defined protein band of approximately 200-220 kDa was obtained for both CD81 and CD53 IPs. This interaction seemed quite robust for CD53, since this band was still seen after IP was performed in the presence of Triton X-100 detergent. Although this molecular weight band was also visible in the CD81 IP, it was clearly less strong than the band in the CD53 IP. Unbiased mass spectrometry analysis identified this band as the protein tyrosine phosphatase CD45 (Eric Rubinstein, personal communication). To confirm these results, the cell surface was biotinylated and IPs were performed for various tetraspanins, including CD53, CD45 itself, and a control membrane protein (CD55). Once again, this high molecular weight band was observed to specifically immunoprecipitate with CD53, in contrast to the other analyzed tetraspanins (CD81, CD82, CD151) and CD55 (Figure 1B). These data indicate that CD45 interacts selectively with CD53 and is not a general tetraspanin partner. Because CD45 is known to be vital in the activation of T cells, we next investigated whether there was also a direct interaction between CD53 and CD45 in the T cell line CEM. In line with the findings in B cells, we observed clear co-IP of CD53 with two CD45 isoforms (CD45RO and CD45RA), indicating that this interaction in also present in T cells (Figure 1C). Based on these findings we report CD53 as a novel interaction partner for CD45 in lymphocytes.

**Figure 1.**
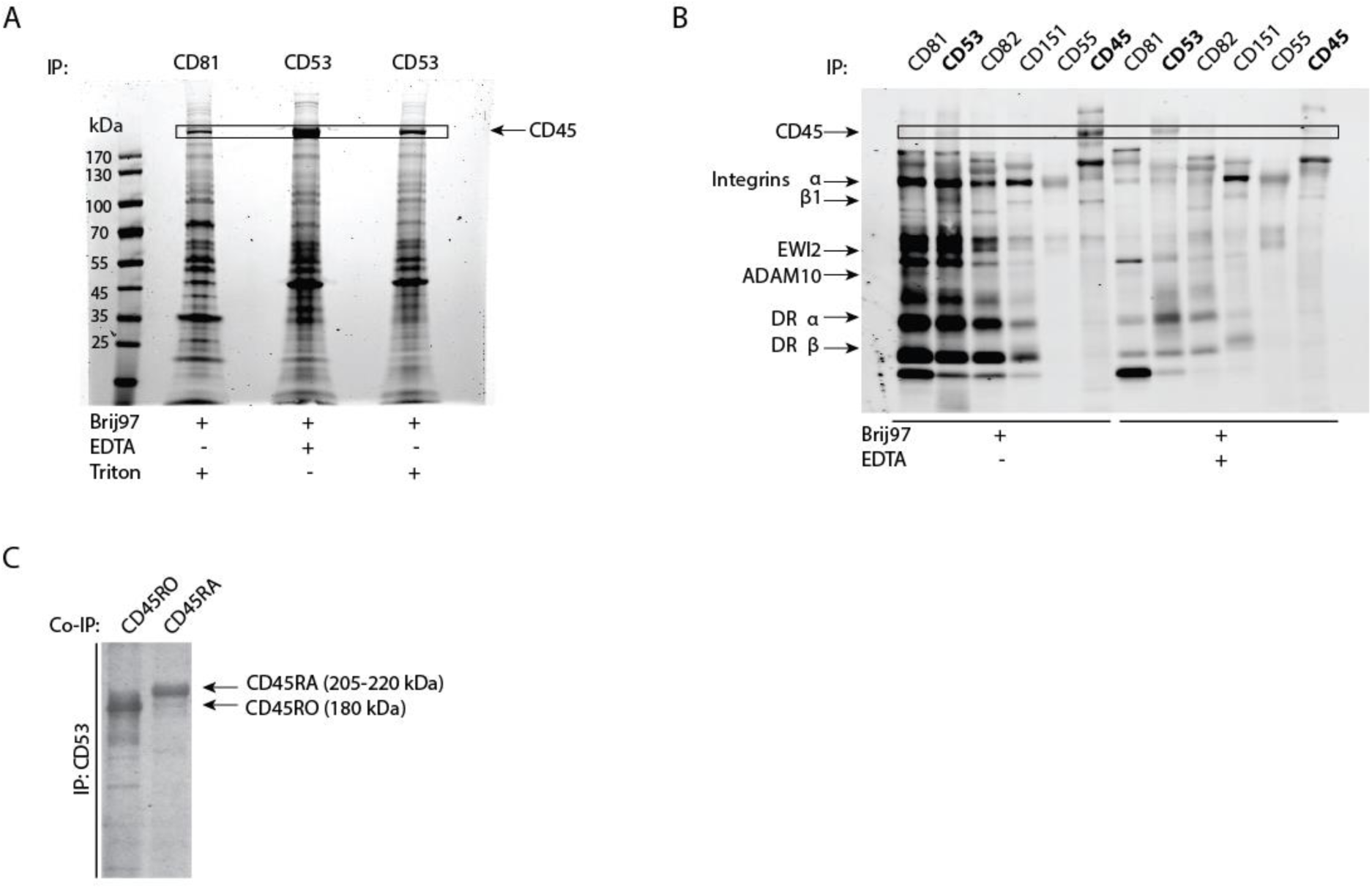
CD45 interacts directly with CD53 in B and T cells. **(A)** SDS-page gel stained with coomassie blue, containing samples from CD53 and CD81 IPs performed in Raji B cells under varying conditions, as indicated at the bottom of the figure. Arrow and outlined box indicate the molecular weight of CD45. **(B)** Western blot of CD81, CD53, CD82, CD151, CD55 and CD45 IPs performed in Raji B cells under varying conditions, as indicated at the bottom of the figure. Cell surface proteins are visualized by biotin labelling. Based on molecular weight, known tetraspanin partners are listed on the left side of the figure, arrows indicate the bands assumed to be attributable to these partner proteins. Outlined box indicates the molecular weight of CD45. **(C)** Western blot of co-IP of CD53 with CD45RO and with CD45RA performed in CEM T cells. Arrows indicate the molecular weight of CD45, CD45RO and CD45RA, respectively. Antibodies directed against CD45RA and RO specifically were used to probe the western blot membrane.

### CD53 acts as co-stimulatory molecule in primary T cell activation

Having established that CD45 specifically interacts with CD53, we hypothesized that CD53 may play a role in activating T cells through CD45. To investigate this, we purified CD4+ and CD8+ T cells from peripheral blood of healthy donors and stimulated these primary human T cells with different combinations of CD3, CD53 and CD28 antibodies and analyzed proliferation in time. Significant proliferation was induced by the combination of anti-CD3 and anti-CD53 (anti-CD3/CD53), or the combination of anti-CD3 and anti-CD28 (anti-CD3/CD28) stimulation in both CD4+ and CD8+ T cell populations (Figure 2A-C). Proliferation induced by the anti-CD3/CD53 combination was considerable higher than that observed for anti-CD3 or anti-CD53 alone, reaching about 50% of the proliferation induced by the positive control (anti-CD3/CD28 stimulation) at 96h in both CD4+ and CD8+ T cells (Figure 2B-C). These data indicate that CD53 can act as co-stimulatory molecule in T cells. In line with this, the anti-CD3/CD53 induced proliferation was characterized by the change from a naïve to an effector T cell phenotype as shown by the switch from CD45RA to CD45RO expression (Figures 2D & E), similar to the anti-CD3/CD28 stimulated T cells. Interestingly, we observed a significantly increased CD53 expression at the T cell surface upon anti-CD3/CD53 or anti-CD3/CD28 stimulation, further implicating CD53 in T cell activation (Figure 2F). Finally, IL-2 production by T cells was readily detected upon anti-CD3/CD28 stimulation, in contrast to anti-CD3/CD53 stimulation (Figure 2G). To analyze whether this was due to retention of IL-2, intracellular IL-2 levels were measured by flow cytometry (Figure 2H). We observed no significant difference in intracellular IL-2 expression between T cells stimulated with anti-CD3/CD53 and cells stimulated with anti-CD3/CD28 indicating that T co-stimulation by CD53 operates via a different pathway than co-stimulation by CD28.

**Figure 2.**
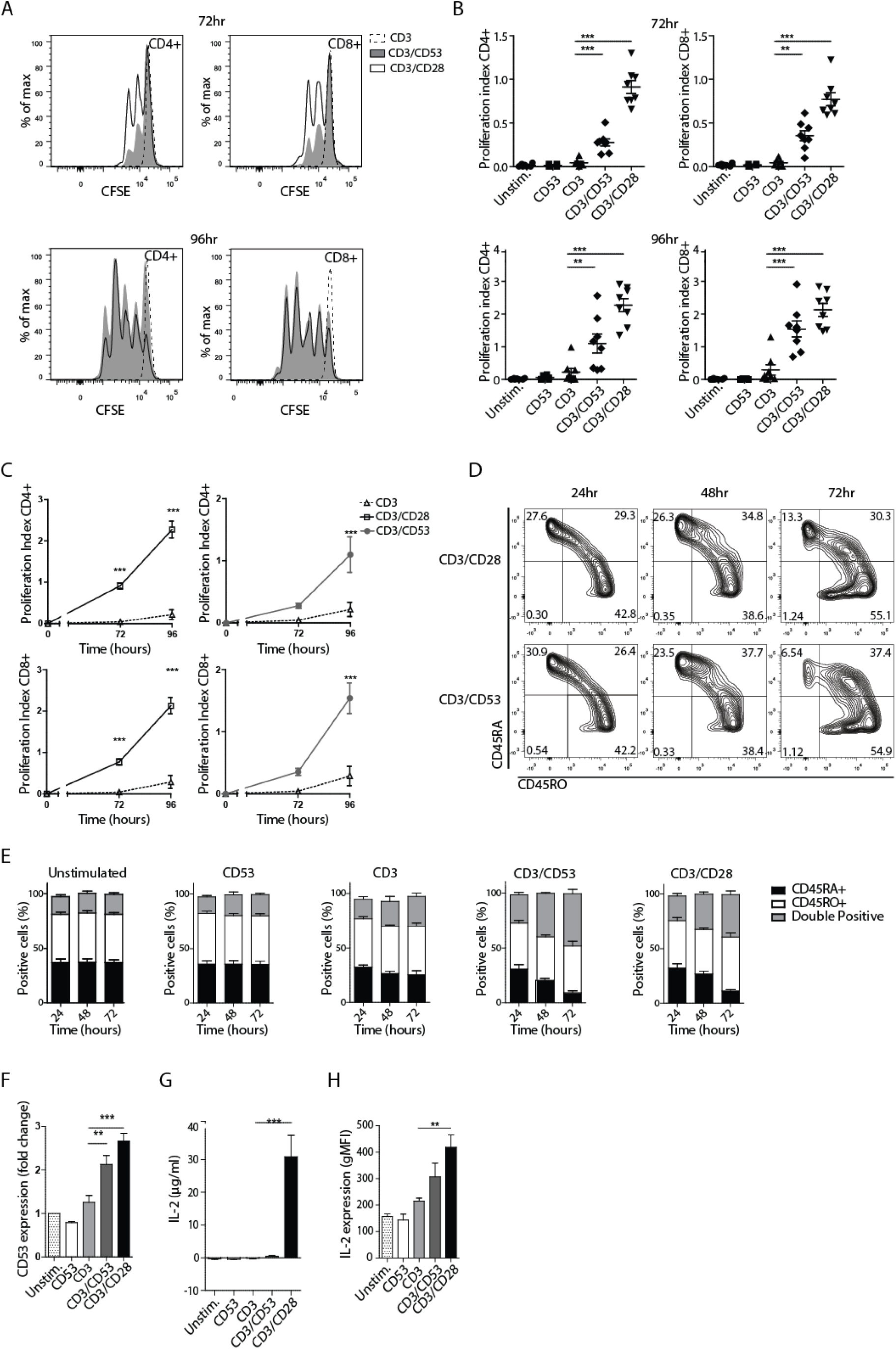
CD53 co-stimulates primary human T cells. **(A)** CFSE measured by flow cytometry for CD4+ and CD8+ T cells of one representative donor at 72 and 96 hours post-stimulation. CD3-, CD3/CD53- and CD3/CD28-antibody stimulated samples are indicated by the dashed line, gray filling and black line, respectively. **(B)** Proliferation index for unstimulated and CD53-, CD3-, CD3/CD53- and CD3/CD28-antibody stimulated CD4+ and CD8+ T cells at 72 and 96 hours post-stimulation. Data points are derived from eight donors collected over four individual experiments. Statistical significance was assessed by unpaired t test. **(C)** Alternate quantification of data presented in (B). Statistical significance was assessed by two-way ANOVA. **(D)** Two-parameter zebra plots depicting the transition of T cells from CD45RA positive to CD45RO positive in time, after stimulation with anti-CD3/CD53 or anti-CD3/CD28, data is derived from one representative donor. **(E)** Quantification of data presented in (D) for six donors over three individual experiments. **(F)** Expression of CD53 on T cells measured by flow cytometry 72 hours after stimulation with either nothing or CD53-, CD3-, CD3/CD53- and CD3/CD28-antibodies. Changes are indicated as fold-change relative to unstimulated samples. Statistical significance was assessed by unpaired t test. Data is derived from six donors over three individual experiments. **(G)** IL-2 production measured by ELISA in cells stimulated for 24 hours with either nothing or CD53-, CD3-, CD3/CD53- and CD3/CD28-antibodies. Statistical significance was assessed by unpaired t test. Data is derived from six donors over three individual experiments. **(H)** IL-2 production measured by intracellular flow cytometry in cells stimulated from 24 hours with either nothing or CD53-, CD3-, CD3/CD53- and CD3/CD28-antibodies. Statistical significance was assessed by one-sample t test. Data is derived from six donors over three individual experiments. All data are means ± SEM (* p<0.05, ** p>0.01, *** p> 0.001)

### T cell proliferation is impaired in the absence of Cd53, but development is normal

In order to better understand the role of CD53 in T cell function, we first investigated whether the absence of CD53 had any effect on the development of T cells. Thymi, spleens and lymphnodes were isolated from 6-week-old *Cd53-/-* and WT (*Cd53+/+* littermate) mice and analyzed for percentages of CD4+ and CD8+ T cell populations. No differences in the percentages of CD4+, CD8+, or double positive T cell populations were observed (Figure S1A), nor was there any difference in the percentage of memory or naïve T cells, in either spleen or lymph nodes between WT and *Cd53-/-* mice (Figure S1B). In addition, similar expression level and percentages of CD3+, CD4+, CD8+ and CD28+ T cells were found in WT and *Cd53-/-* mice, with the exception of the percentage of CD3+ cells in the lymph node compartment which was higher in *Cd53-/-* mice (Figure S1C) due to a B cell deficit (Mark Wright, personal communication). Taken together, these data indicate that CD53 is not required for T cell development in mice. Next, we investigated whether the absence of CD53 altered the functional capacity of these cells. Splenic T cells were isolated from WT and *Cd53-/-* mice and labeled to track cell division upon anti-CD3/CD28 stimulation. We observed that proliferation was markedly reduced for both CD4+ and CD8+ *Cd53-/-* T cells compared to WT T cells (Figure 3A-B). Quantification revealed a significantly decreased proliferation index for *Cd53-/-* T cells compared to WT T cells for both CD4+ and CD8+ subsets (Figure 3C & D). Additionally, further characterization of these T cells revealed no differences in the expression of the activation markers CD25 and CD69 on the surface of WT and *Cd53-/-* T cells after stimulation with PMA or anti-CD3/CD28 (Figure 3E). Moreover, no differences were seen in the levels of IL-2 produced by these cells after anti-CD3/CD28 stimulation (Figure 3F). Overall, these results indicate that the activation of T cells is impaired in the absence of CD53, and that the role of CD53 in this activation is independent of IL-2 production.

**Figure 3.**
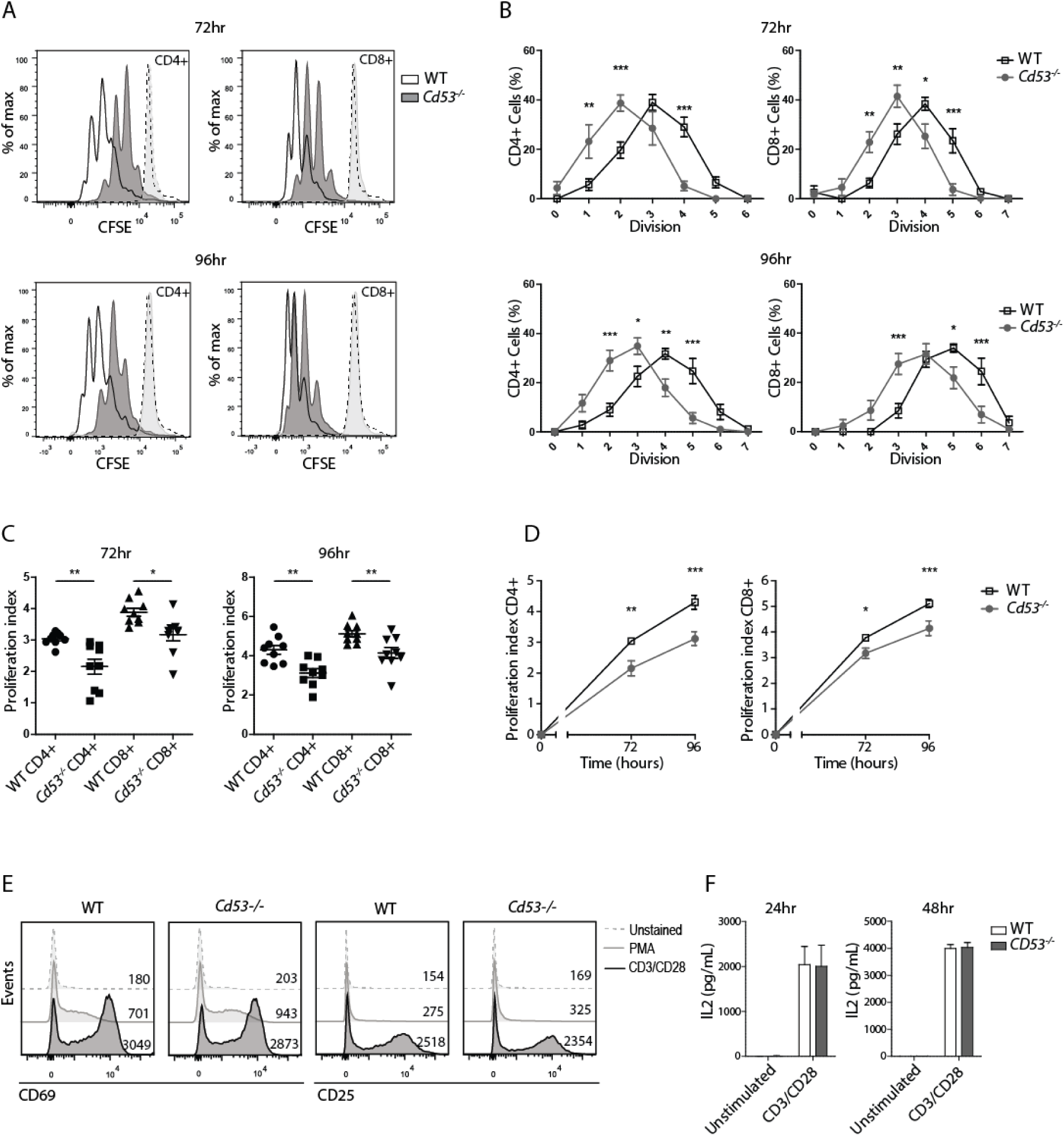
*Cd53-/-* mice show impaired T cell proliferation responses. **(A)** CFSE measured by flow cytometry for CD4+ and CD8+ T cells of one WT and one *Cd53-/-* mouse at 72 and 96 hours post-stimulation. Anti-CD3 stimulated samples are indicated by dashed line and light gray for WT and *Cd53-/-*, respectively. Anti-CD3/CD28 stimulted samples are indicated by solid black line and dark gray for WT and *Cd53-/-* respectively. **(B)** Percentage of CD4+ and CD8+ T cells per division as tracked by CFSE at 72 and 96 hours post-stimulation. WT and *Cd53-/-* data points indicated by unfilled squares and gray filled circles, respectively. Data is derived from nine mice per genotype, collected over three individual experiments. Statistical significance was assessed by two-way ANOVA. **(C)** Proliferation index of WT and *Cd53-/-* T cells at 72 and 96 hours post-stimulation, CD4+ and CD8+ T cell populations shown separately. Data points are derived from nine mice per genotype collected over three individual experiments. Statistical significance was assessed by unpaired t test. **(D)** Alternate quantification of data presented in (C). Statistical significance was assessed by two-way ANOVA. **(E)** Expression of CD69 and CD25 on WT and *Cd53-/-* T cells as measured by flow cytometry 24 hours after stimulation with either nothing, PMA or anti-CD3/CD28. gMFI of signal is indicated per graph on the right side. Data was gathered from six mice per genotype from two independent experiment. Representative data from WT and *Cd53-/-* mice is shown. **(F)** IL-2 production measured by ELISA for WT mice (white bars) and *Cd53-/-* mice (gray bars) at 24 and 48 hours after stimulation with either nothing or anti-CD3/CD28. Data is derived from nine to ten mice per genotype per time point, collected over five independent experiments. Statistical significance was assessed by unpaired t test. All data are means ± SEM (* p<0.05, ** p>0.01, ***p> 0.001).

### Cd53-/-T cells have impaired recall capacity in vivo

Based on our finding that naïve CD53-negative T cells are impaired in proliferation, we decided to investigate whether this also extended to T cell recall responses based on *in vivo* activation. To analyze the function of CD53 on T cells *in vivo*, we investigated T cell recall responses by immunizing *Cd53-/-* and WT mice with multiple doses of keyhole limpet haemocyanin (KLH) adsorbed to the adjuvant alum, or immunization with PBS as a control (Figure 4A). Post-immunization, isolated T cells were labeled to track proliferation and restimulated with KLH in the presence of naïve WT APCs. In line with our *in vitro* data, we observed that the *in vivo* primed, antigen-specific, CD4+ T cells from *Cd53-/-* mice, showed reduced proliferation upon restimulation with KLH compared to WT T cells (Figure 4B). Non immunized (PBS injected) mice showed no significant proliferation for either genotype, ensuring that the response measured was specifically induced by recognition of the KLH antigen. Quantification revealed significantly lower proliferation indexes for the CD4+ T cells of *Cd53-/-* mice compared to the WT mice (Figure 4C-D). Moreover, a significantly higher percentage of T cells from WT mice entered cell division compared to T cells from *Cd53-/-* mice. Taken together, these results demonstrate that the absence of CD53 impairs the T cell response *in vivo*.

**Figure 4.**
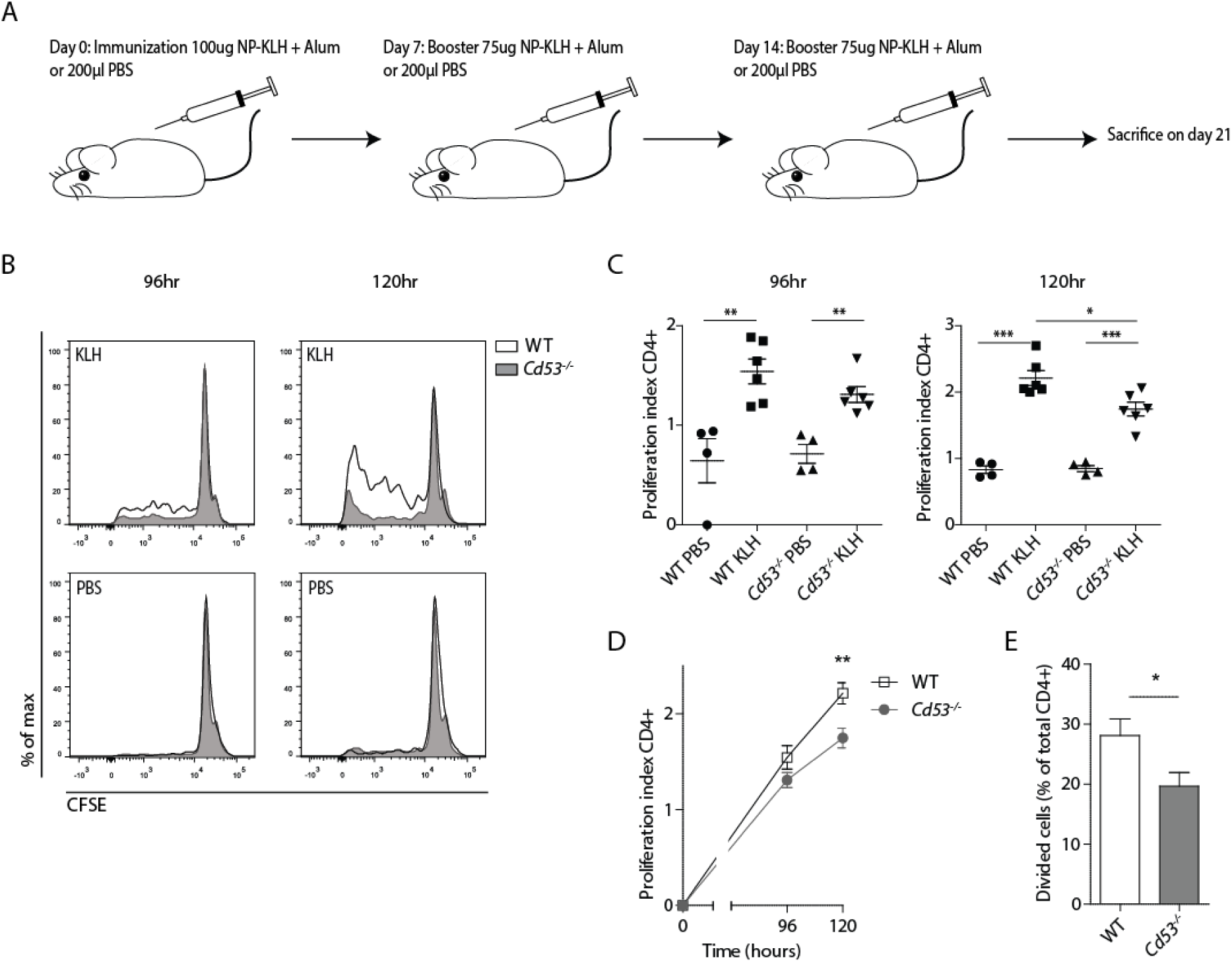
T cells of Cd53-/-mice show impaired recall response post-immunization. **(A)** Graphical overview of experimental schedule and design. **(B)** CFSE measured by flow cytometry for CD4+ T cells of both KLH and PBS immunized WT mice (black line) and *Cd53-/-* mice (gray filled) at 96 and 120 hours post-stimulation with 100 μg/ml KLH in the presence of naive WT APCs. Data was gathered from six mice per genotype from two independent experiments. Representative data from one WT and one *Cd53-/-* mouse is shown. **(C)** Proliferation index of CD4+ T cells from PBS and KLH immunized WT and *Cd53-/-* at 96 and 120 hours post-stimulation. Data points are derived from six mice per genotype collected over two individual experiments. Statistical significance was assessed by unpaired t test. **(D)** Alternate quantification of data presented in (C). Statistical significance was assessed by two-way ANOVA. **(E)** Percentage of cells from the total CD4+ population, isolated from KLH immunized mice, entering division upon KLH restimulation in vitro. Data points are derived from six mice per genotype collected over two indepenent experiments. Statistical significance was assessed by unpaired t test. All data are means ± SEM (* p<0.05, ** p>0.01, *** p> 0.001).

### CD53 is recruited to the site of TCR activation but does not interact with PKCθ

Since CD45 is known to function during early TCR signaling at the immunological synapse, we investigated whether CD53 was also recruited to sites of heavy TCR signaling. To achieve this, we examined the localization of CD53 during TCR signaling using the micro-contact printing technique_14_. This allows us to analyze the recruitment of membrane proteins to discrete areas of TCR signaling in live cells. Jurkat T cells were transfected with CD53-green fluorescent protein (GFP) or CD37-GFP as control. Transfected T cells were seeded on surfaces containing either anti-CD3/CD28 or isotype control antibodies, after which localization of CD53 and CD37 was assessed. We observed specific recruitment of CD53 to the contact site with the anti-CD3/CD28 stamp, but not to the isotype stamp (Figure 5A). CD37 on the other hand showed no recruitment to either the isotype or the anti-CD3/CD28 stamp, indicating that the recruitment triggered by TCR activation is specific to CD53 (Figure 5B). Quantification showed that the CD53 signal was significantly increased at the stamp contact site, in contrast to CD37 (Figure 5C). Based on our previous study that demonstrated a direct interaction between CD53 and PKCβ in B cells, we investigated whether such interaction was also present in T cells_14_. Using micro-contact printing we found that PKCθ was also recruited to the TCR signaling sites, which was contemporaneous with CD53 recruitment (Figure S3A-B). Next, we assessed whether there was a direct interaction between CD53 and PKCθ using fluorescence imaging microscopy (FLIM). Jurkat T cells transfected with either CD53-mCitrine alone or with CD53-mCitrine and mCherry-PKCθ were stimulated with phorbol myristate acetate (PMA) to induce PKCθ membrane recruitment. We observed no decrease in the lifetime of donor fluorophore mCitrine upon translocation of PKCθ to the membrane, indicating that there is no direct interaction between CD53 and PKCθ upon T cell activation (Figure S3C). Together these data demonstrate that tetraspanin CD53 is selectively recruited to TCR signaling sites. In addition, CD53 does not control the recruitment and/or stabilization of PKC upon T cell stimulation.

**Figure 5.**
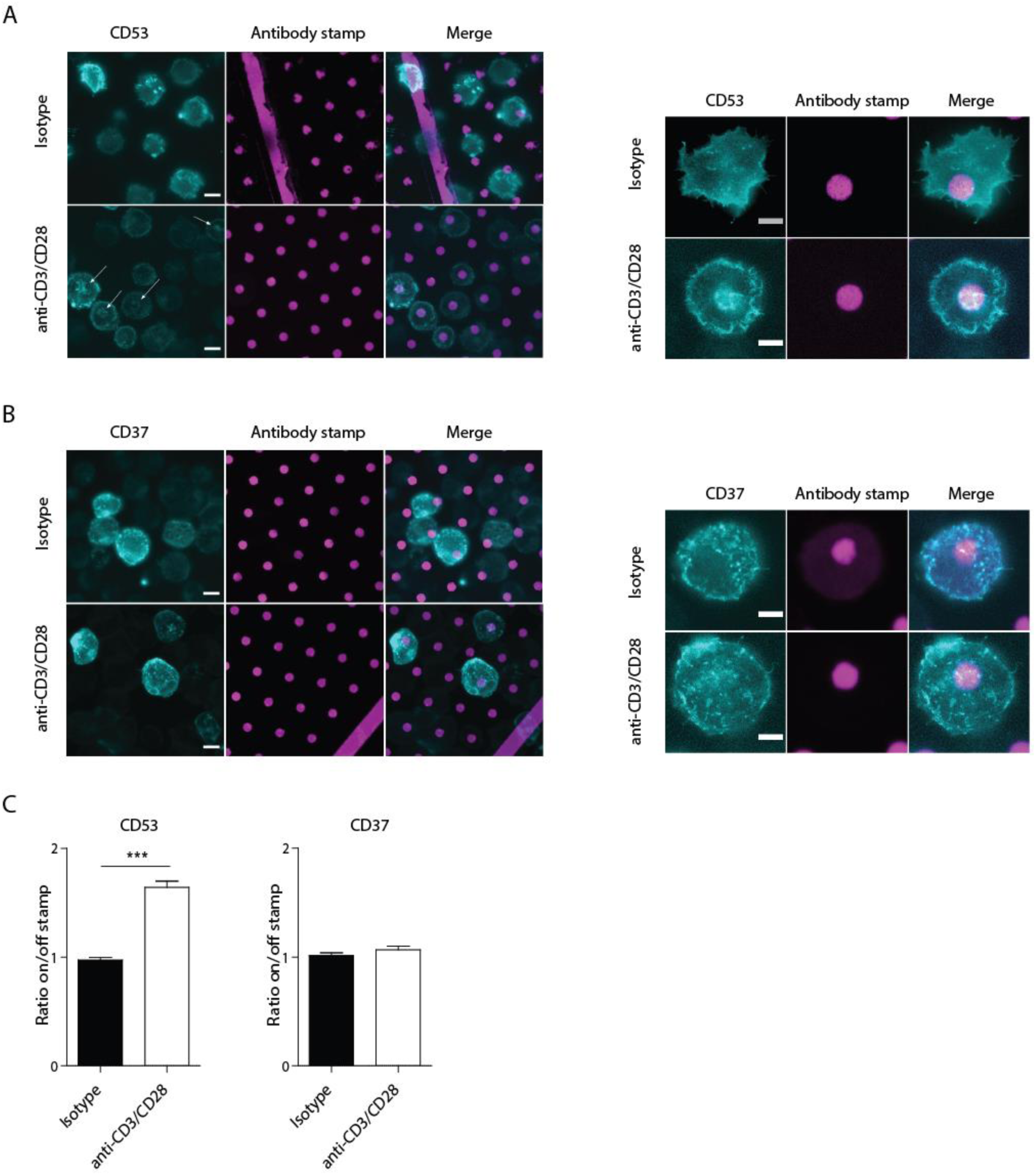
CD53 is recruited to TCR signaling sites. **(A)** Jurkat T cells expressing GFP-CD53 were seeded on isotype control or anti-CD3/CD28 prints and CD53 recruitment was analyzed after 8 minutes by epifluorescence microscopy. Left: overview of multiple cells. Right: larger magnification of one representative cell. **(B)** Jurkat T cells expressing GFP-CD37 were seeded on isotype control or anti-CD3/CD28 prints and analyzed after 8 minutes by epufluorescence microscopy. Left: overview of multiple cells. Right: larger magnification of one representative cell. **(C)** Ratio between the average CD53 (left) or CD37 (right) intensity on the antibody print and that outside of the print. Data represents analysis of >30 cells from three independent experiments. Statistical significance was assessed by one-sample t test. Data are means ± SEM (* p<0.05, ** p>0.01, *** p> 0.001). Scale bars represent either 10 μm (left panels) or 5 μm (right panels) as indicated in the bottom right corner of the leftmost images. Arrows indicate TCR signaling sites showing CD53 recruitment.

### CD53 is important for the mobility and stability of CD45RO on the T cell surface

Our findings indicate that tetraspanin CD53 interacts specifically with CD45 in T cells. To better understand this interaction, we created a human *CD53-/-* T cell line by applying CRISPR/Cas9 technology (Figure 6A). General characterization of the *CD53-/-* T cell line revealed no significant differences in the expression of CD3, CD4 and CD28 protein compared to the WT parental cell line (Figure S2). Interestingly, upon loss of CD53, T cells exhibited a striking change in the expression profile of CD45. Despite observing only, a slight overall decrease in the total expression of CD45, a remarkable difference was observed in the expression of CD45RA and CD45RO between *CD53-/-* and WT T cells (Figure 6B). WT T cells expressed predominantly CD45RO (approx. 60%), with a smaller population (approx. 20%) of CD45RA-positive cells. This profile was inverted in *CD53-/-* T cells, with a small CD45RO-positive population (approx. 20%), and a large CD45RA-positive population (approx. 50-60%) present in these cells. This remarkable difference in CD45 isoform expression induced only by the removal of CD53 from the cell surface, led us to hypothesize that CD53 may be specifically required for the stabilization of CD45RO at the plasma membrane. In order to investigate whether CD53 interacts with CD45RO in order to stabilize this protein, fluorescence recovery after photo-bleach (FRAP) was applied. *CD53-/-* and WT cells transfected with CD45RO fused to GFP were subjected to FRAP localization of CD45RO (Figures 6C-F). These findings combined with the loss of CD45RO expression on the surface of *CD53*-negative T cells, may be explained by a reduced stability of CD45RO in the absence of CD53. To microscopy. We observed an increased mobility of CD45RO in *CD53-/-* T cells compared to WT T cells, indicating that CD53 does interact with CD45RO, in turn regulating the spatio-temporal investigate this, CD45 internalization analysis was performed on WT and *CD53-/-* T cells. The results demonstrated that CD45RO had a significantly higher rate of internalization in the absence of CD53 (74.53% CD45RO positive cells in WT versus 52.57% in KO after 24h). This finding confirmed that the stability of CD45RO on the T cell membrane is dependent on tetraspanin CD53 (Figure 6F-G). Together our findings demonstrate that CD53 is important for the mobility and stability of CD45RO on the T cell surface.

**Figure 6.**
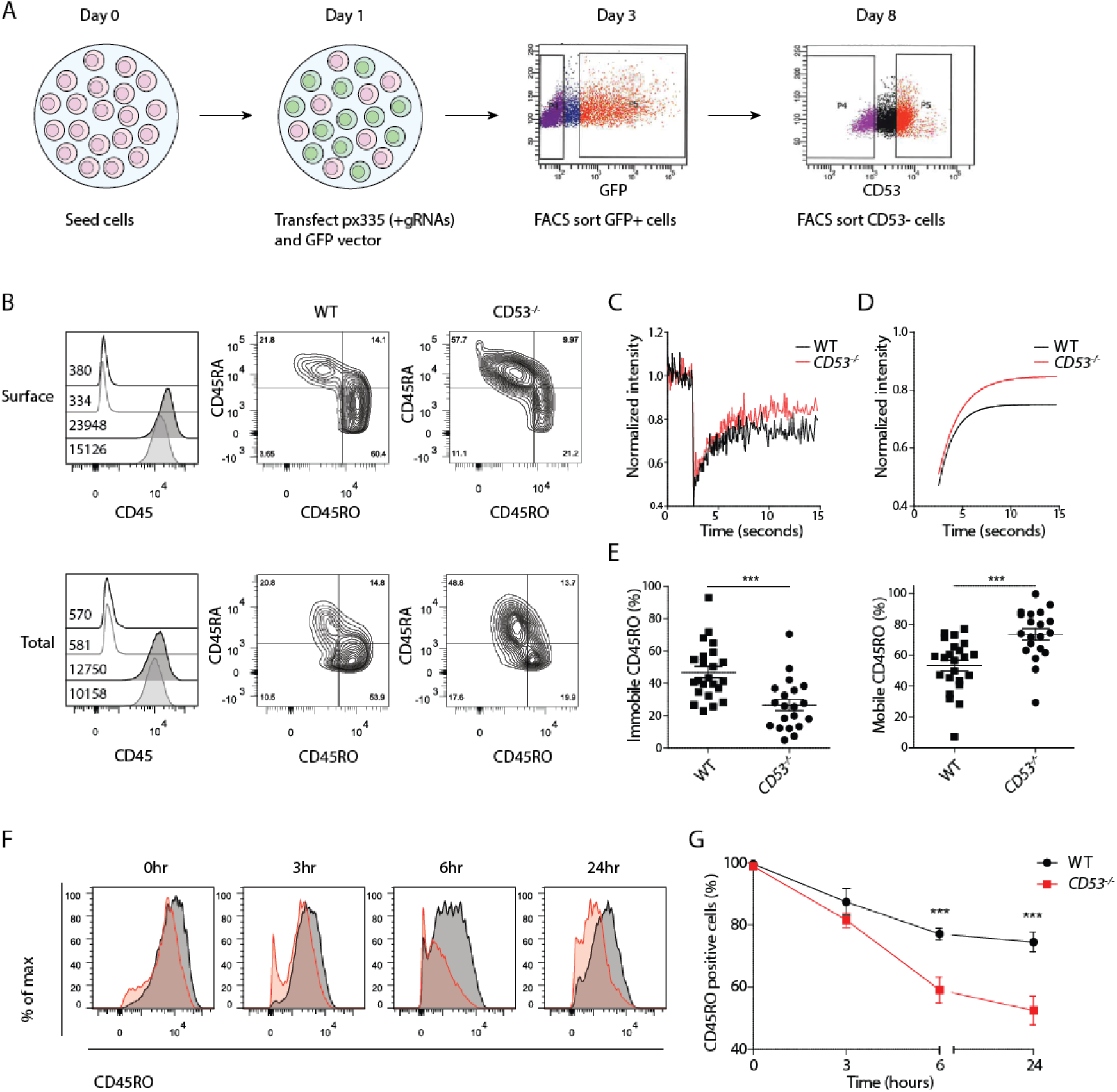
CD53-deficiency results in defective CD45 expression and increased mobility of CD45RO at the T cell surface. **(A)** Graphical overview of the CRISPR/Cas9 technique as applied to CEM T cells to knock out CD53. **(B)** Left: surface and total expression of CD45 in WT and *CD53-/-* CEM T cells as measured by flow cytometry. Unfilled black and grey lines reprepresent unstained WT and *CD53-/-* CEM T cells, respectively. Black and grey filled graphs represent pan-CD45 stained WT and *CD53-/-* CEM T cells, respectively. Middle and right: two-prameter zebra plot showing the surface and total expression of CD45RA and CD45RO for WT and *CD53-/-* CEM T cells. Data was gathered from three independent experiments, representative data from one experiment is shown. **(C)** FRAP curve for CD45RO in WT and *CD53-/-* CEM T cells, curve represents the average of all cells from one representative experiment of three independent experiments. Curve is based on >10 cells per genotype. **(D)** Fit of the curve obtained in (C) as calculated by OriginPro_8_ softare using formula for bi-exponential decay. **(E)** Percentage of immobile (left) and mobile (right) CD45RO as calculated based on FRAP curves. Data points are derived from >20 cells per genotype collected over three independent experiments. Statistical significance was assessed by one-sample t test. **(F)** Internalization of CD45RO over time as measured by flow cytometry. Filled black and red graphs denote WT and *CD53-/-* CEM T cells, respectively from one representative experiment. **(G)** Percentage of CD45RO-positive cells in time based on quantification of CD45RO internalization as shown in. **(F)** Data is derived from three independent experiments. Statistical significance was assessed by two-way ANOVA. All data are means ± SEM (* p<0.05, ** p>0.01, *** p> 0.001).

## Discussion

The protein tyrosine phosphatase CD45 is essential for T cell proliferation, yet it remains poorly understood how CD45 isoforms are regulated_27, 28_. Here we report that tetraspanin CD53 is required for CD45RO expression and stabilization at the T cell surface, which is of immunological importance as evidenced by the impaired TCR activation and proliferation of CD53-deficient T cells. This novel interaction provides insight into both the (isoform-specific) regulation of CD45 as well as the role of CD53 in T cell function. To the best of our knowledge, this is the first identification of a membrane partner that directly regulates the stabilization and mobility of a specific CD45 isoform on the T cell surface.

Other proposed CD45-interacting proteins on the T cell surface include, CD45-AP, CD4/CD8 and CD2. The absence of CD45-AP, which specifically associates with CD45, has been found to affect the interaction between CD45 and Lck suggesting that CD45-AP either directly or indirectly mediates this association_29_. In *Cd45-AP-/-* mice, this resulted in an impaired T cell proliferation, similar to what we have observed in T cells derived from *Cd53-/-* mice_29_. Given these comparable findings, it would be interesting to investigate whether CD53 is a part of the reported supramolecular protein complex formed by CD45-AP, CD45, CD4 and Lck_30_. Interestingly, the IPs we performed in T cells identified the CD45RO isoform as a specific partner of CD53. Similarly, CD4 and CD8 have been shown to interact with CD45RO, but not with the CD45RBC or CD45RABC isoforms_31_. This interaction was found to have functional implications, as the expression of CD45RO was associated with enhanced activity of CD4-associated Lck. The fact that both CD53 and CD4/CD8 preferentially interact with CD45RO, provides an interesting possible explanation for the observed proliferation defect which merits further investigation. In addition, the expression of CD45-AP has been found to depend on the presence of CD45 on the membrane. Since we have found that total CD45 expression is affected by CD53, the subsequent loss of CD45-AP is an additional possible mechanism by which CD53 could regulate T cell function. Lastly, CD2 has been suggested to be functionally associated with CD45 through crosslinking experiments_32_. In line with this, antibodies directed against CD45 have been shown to induce strong T cell proliferation when combined with anti-CD2 antibodies, but not when combined with anti-CD3 antibodies. CD53 has also been shown to associate with CD2 in rat T cells, suggesting that CD53 may act as a possible intermediary protein, connecting CD45 and CD2, thereby modulating proliferative responses of T cells.

In addition to identifying this novel association, we have also provided evidence that the interaction between CD53 and CD45RO is important for the spatial regulation and stabilization of CD45RO on the T cell membrane. This in turn affects the overall isoform expression pattern of CD45. This is notable since recent evidence has shown that CD45 function is also dependent on combinations of CD45 isoforms expressed_33_. A striking change in the isoform expression profile of CD45 was observed when CD53 was removed from the cell surface, with CD45RO expression being severely reduced in CD53-negative cells while CD45RA expression increased (Figure 6B). Our findings showed that in the absence of CD53, CD45RO was significantly less stable on the cell surface and showed an enhanced rate of internalization compared to WT T cells (Figure 6F-G). It remains to be seen whether primary CD53-negative T cells only exhibit reduced CD45RO expression, or whether they also show (re-)expression of alternative CD45 isoforms. Supporting this, we observed re-expression of CD45RA upon CRISPR-knockout of CD53 in human T cells. Similarly, re-expression of longer CD45 isoforms has been detected in human memory T cells_34_. CD45 isoform switching occurs naturally during the course of normal T cell activation, yet despite much effort, the exact purpose of this process remains largely unclear. This is believed to be important though, since the alternative splicing of CD45 is highly regulated, and conserved in vertebrate evolution_35, 36_. In line with this, some studies have shown isoform dependent differences in downstream signaling, though the interpretation of these results is sometimes complicated by seemingly contradictory findings_37_. For example, CD45RO has been reported to preferentially associate with CD4 and CD8, leading to increased signaling via Lck in T cells expressing higher levels of CD45RO compared to other isoforms_31_. Additionally, others have studied the ability of CD45 isoforms to participate in enhancing T cell activation, with CD45RO found to be most effective at this_38_. Other studies have presented seemingly contradictory results, showing that CD45RO was less capable of inducing proliferation in thymic T cell compared to the full length CD45RABC, though this may be affected by the developmental stage of these cells_39_. More recently, using a model membrane system, it has been shown that isoforms of CD45 can segregate differently during TCR-pMHC interaction, suggesting this may be a mechanism for the fine-tuning of signaling_40_. With so much uncertainty still surrounding the role of specific isoforms of CD45 in T cell biology, our findings contribute to a better understanding of this by providing evidence of an isoform-specific interaction with CD53 which has clear functional consequences.

As a membrane organizing protein, CD53 likely contributes to the spatio-temporal regulation of CD45RO, of which very little is known_41, 42, 43_. The importance of this spatio-temporal organization is illustrated by studies in the immunological synapse (IS) in which the dynamic inclusion and exclusion of CD45 is known to be important for balancing the opposing roles of CD45 in T cell signaling. As a result, CD45 can function as both a positive and a negative factor in T cell activation, depending on its spatio-temporal organization and local concentration_7, 44_. In the past, numerous mechanisms have been proposed to regulate the activity of CD45, these include localization and dimerization_36_. CD53 exerts its function through the formation of tetraspanin enriched microdomains_45_. The regulation of CD45 by membrane microdomains has been suggested before, with findings indicating that lipid raft-associated CD45 acts as a negative regulator of TCR signaling, while CD45 located outside of lipid rafts showed minimized opposition to TCR signaling_46_. Studies using purified detergent-resistant membranes (DRMs) should be interpreted with caution as DRMs not only contain lipid rafts but also tetraspanin enriched microdomains _47_. Additionally, passive regulation of CD45 localization during T cell activation has been proposed by the kinetic segregation model, which suggests that the tight contact zones in which the TCR and MHC interact exclude large molecules such as CD45 based on size_48_. CD45 chimeras, with ectodomains derived from other proteins, showed that smaller ectodomains blunted the IL-2 response upon T cell activation_48_. Though the authors propose that this argues in favor of the kinetic segregation model, ectodomain exchange would also clearly affect interaction with partner molecules and the ability to dimerize, a major issue not taken into account in these studies. Given our finding that CD53 accumulates at sites of TCR signaling, it is possible that CD53 may be contributing in some way to the organization or localization of CD45RO at the immune synapse. This does not refute or support the theory of kinetic segregation, as these could exist as complementary processes during T cell activation. Furthermore, cytoskeletal contacts through proteins like spectrin and ankyrin have also been proposed to participate in regulating the localization of CD45, with disruption of the actin-spectrin interaction leading to increased CD45 lateral mobility, similar to what we have observed for the mobility of CD45RO in the absence of CD5349.

The dimerization of CD45 has been suggested to negatively regulate CD45 activity, with the RO isoform dimerizing most efficiently_50_. For other receptor-like protein tyrosine phosphatases (RPTPs) dimerization has been established as a regulatory mechanism, but for CD45 this remains somewhat speculative though the wedge structure involved in dimerization of other RPTPs is conserved in CD45_51, 52_. Supporting a role for dimerization, mutation of the wedge structure in CD45 molecules seemingly promotes signaling activity_53_. Additionally, a chimeric version of CD45 containing the ligand binding domain of epidermal growth factor receptor (EGFR) lost the capacity to support TCR signaling upon induced dimerization. In contrast, it has been proposed that based on the crystal structure of the cytoplasmic D1-D2 segment of CD45, dimerization would not be possible according to the wedge model proposed_54_. Alternatively, there is evidence to support the involvement of the extracellular domain in homodimerization of CD45RO_31_. Although the exact role that dimerization plays in the regulation of CD45 cannot be fully established, we cannot exclude that CD53 may be involved in modulating CD45 dimerization though specific organization.

Based on our findings we propose a model which illustrates the role of CD53 in the regulation of CD45RO (Figure 7). We posit that the interaction between CD53 and CD45RO is required in order for T cells to effectively undergo CD45 isoform switch upon activation. In the absence of CD53, the expression of CD45RO is deregulated, leading to a disturbed or incomplete T cell activation, which results in an impaired proliferation. We hypothesize that in the absence of CD53 the expression of CD45RO is reduced due to membrane instability of this isoform. The impaired expression of CD45RO on the cell surface can have a dual effect, related to both the loss of CD45RO-specific signaling as well as the alteration of the total CD45 isoform expression profile, both of which can contribute to an altered T cell signaling. This effect, coupled with the increased mobility of any remaining surface CD45RO, contribute to a diminished TCR signaling capacity leading to the reduced proliferation observed for CD53-negative T cells.

**Figure 7.**
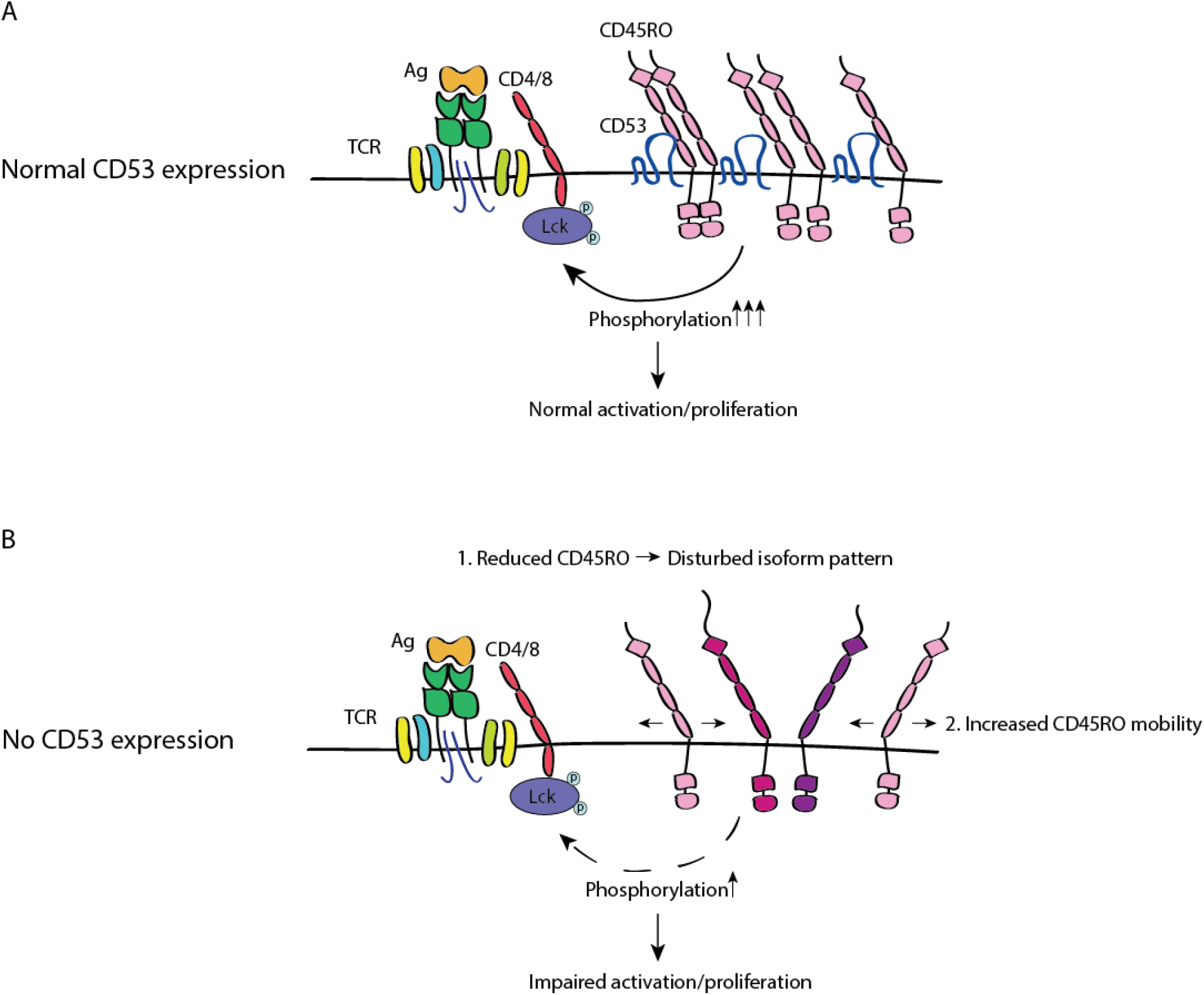
CD45RO surface stability depends on CD53. Model depicting the proposed role of CD53 as a regulator of CD45RO at the cell surface. **(A)** In the presence of CD53 the stability of CD45RO is enhanced, and mobility is reduced. Together these two factors enhance CD45 function and potentiate phosphorylation downstream of the TCR leading to normal levels of activation and proliferation. **(B)** In the absence of CD53, CD45RO is less stable leading to reduced expression. This produces an altered CD45 isoform expression profile, different isoforms of CD45 are indicated by varrying colors. Both of these outcomes have consequences for the signaling capacity of the TCR. In addition, the CD45RO remaining on the cell surface shows enhance mobilty, which can also contribute to the reduced efficiency of CD45 in CD53-negative T cells resulting in diminished activation and proliferation.

Our findings not only shed light on the biology of CD45, but also contribute significantly to a better understanding of CD53 function in the immune system. In this study, we measured the proliferative capacity of WT and *Cd53-/-* primary murine T cells both *in vitro* and *in vivo*. Our findings demonstrate that CD53 is important for T cell activation, as CD53-negative T cells were significantly impaired in their proliferative capacity (Figures 3 and 4). The importance of CD53 was confirmed in primary human T cells, which could be co-stimulated through cross-linking of CD53 (Figure 2). In line with our findings, antibody cross-linking of tetraspanins including CD82, CD81, CD9 and CD63, has previously been shown to have a co-stimulatory effect on T cells_55, 56, 57, 58_. Deficiency of Tssc6, CD37, CD81 and CD151 have all been linked to hyper-proliferation of primary murine T cell_59, 60, 61, 62_. This is in stark contrast to our findings, which showed a reduced proliferation of *Cd53-/-* T cells, suggesting that CD53 on T cells functions through a mechanism that is unique to this tetraspanin. Furthermore, we have previously reported that CD53 plays an important role in recruiting and stabilizing PKCβ in B cells_14_. In contrast, our observations show that the CD53 is not involved in the direct stabilization of PKCθ in T cells. This illustrates the versatility of CD53 as a regulator of signaling, exhibiting multiple modes of interaction with the lymphocyte activating pathways controlling T and B cell responses.

In conclusion, we demonstrate that there is isoform-specific regulation of the expression of CD45RO on the T cell surface by tetraspanin CD53. This interaction is shown to have functional consequences, as the absence of CD53 negatively affects proliferation of T cells, while CD53 cross-linking functions as a co-stimulatory signal. Our findings shed new light on the molecular mechanisms by which CD45 is regulated, and places CD53 in a unique position, as a direct and specific regulator of CD45RO in T cells. Perhaps most excitingly, these findings present us with a potential new means by which to modulate T cell activity in immune related diseases.

## Materials and Methods

### Antibodies and dyes

The following antibodies were used: anti-human CD53 (MEM-53, Biorad), anti-human CD53-488 (MEM-53, Novus Biologicals), anti-human CD3 (OKT3, Bio X cell, anti-human CD28 (9.3 Bio X Cell), anti-human CD45 (REA747, Miltenyi), anti-human CD45RA-PE (HI30,Biolegend), anti-human CD45RO-APC (UCHL1, BD Pharmingen), anti-human CD3-PE (HIT3a, BD Pharmingen), anti-human CD4-APC (RPA-T4, Biolegend), anti-human CD4-PerCP (RPA-T4, Biolegend), anti-human CD8-FITC (RPA-T8, BD Pharmingen), anti-human CD8-APC (RPA-T8, BD Pharmingen), anti-human CD25-PE (M-A251, BD Pharmingen), anti-human CD69-PerCP (L78, BD Pharmingen), anti-human CD53 (TS5315), anti-human CD45 (IOL1, Immunotech), anti-human CD45RA (MEM-56, Novus Biologicals), anti-human CD45RO (UCHL-1, Thermo Fisher Scientific), anti-mouse CD3 (17A2, Bio X Cell), anti-mouse CD28 (37.51, Bio X Cell), anti-mouse CD28 (37.51, BD Pharmingen), anti-mouse CD3-APC (145-2C11, Biolegend), anti-mouse CD3-FITC (17A2, Biolegend), anti-mouse CD4-FITC (RMA4-5, Biolegend), anti-mouse CD4-PerCP (RMA4-5, Biolegend), CD8-PE (53.6-7, BD), CD8-PerCP (53.6-7, BD Pharmingen), anti-mouse CD25-APC (PC61.5, eBioscience), anti-mouse CD69-PE (H1.2F3, BD Pharmingen), anti-mouse CD44-PeCy7 (IM7, Biolegend), anti-mouse CD62L-BV510 (MEL-14, Biolegend), goat anti-Syrian hamster IgG H&L Alexa Fluor 647 (Invitrogen), anti-mouse IL2-PE (JES6-5H4, Biolegend), anti-mouse IL-2 (JES6-1A12, BD), anti-mouse IL-2 biotin (JES6-5H4, BD),CellTrace Violet (Thermo Fisher Scientific).

### Cell lines

Human T cell lines, CCRF-CEM and Jurkat, and the human B cell line Raji were used which are all widely used to study lymphocyte activation and signaling 16, 17. Jurkat, CEM and Raji cells were maintained at 37 °C in 5 % CO2. Cell were cultured in RPMI 1640 (Gibco) supplemented with 10 % fetal bovine serum (FBS; Hyclone), 1 % stable glutamine (PAA) and 1 % antibiotic-antimycotic (Gibco).

### Isolation of human primary T cells

Human primary T cells were isolated from buffy coats obtained from healthy volunteers and in accordance with the recommendations of institutional guidelines. All subjects gave written informed consent in accordance with the Declaration of Helsinki. Pan T cells were isolated from peripheral blood leukocytes using the Pan T cell isolation kit according to manufacturer’s instructions (Miltenyi Biotec). Isolated human T cells were cultured in *X-VIVO*-15 medium supplemented with 2 % human serum.

### Mice

Generation of *Cd53*−/− mice has been described previously (Zuidscherwoude et al. 2017). *Cd53*−/− mice and sex- and aged-matched C57Bl/6J WT (*Cd53+/+*) littermate mice were bred at the Central Animal Laboratory, Nijmegen (The Netherlands). Mice were housed in top-filter cages and fed a standard diet with freely available water and food and used at 8 to 12 weeks of age. All *in vivo* studies complied with European legislation and were approved by the local animal ethical committee for the care and use of animals with related codes of practice.

### Isolation of murine primary T cells

Resting primary immune cells were isolated from either thymus, spleen or inguinal lymph nodes of WT and *Cd53-/-* mice (as indicated in figure legends). Organs were digested using collagenase type III (Worthington) and DNAse I (New England Biolabs) before being passed through a 100 μm filter. T cells were isolated using the Pan T Cell isolation kit II according to the manufacturer’s instructions (Miltenyi Biotec). Cells were cultured in RPMI 1640 (Gibco) supplemented with 10 % fetal bovine serum (FBS; Hyclone), 1 % stable glutamine (PAA) and 1 % antibiotic-antimycotic (Gibco), 1mM sodium pyruvate and 0.1 % 2-mercaptoethanol. Isolated cells were used for experiments as described below.

### Cell labeling and immunoprecipitation

For surface labeling, cells were washed three times in Hank’s buffered saline and incubated in PBS containing 0.5 mg/ml EZ-link-Sulpho-NHS-LC-biotin biotin (Pierce, Rockford, IL, USA). After 30 min of incubation at 4 °C, cells were washed three times in PBS to remove free biotin. Labeled cells were lysed at the concentration of 2 x107/ml in lysis buffer containing 10 mM Tris, pH 7.4, 150 mM NaCl, 0.02% NaN3 and protease inhibitors and supplemented with 1% Brij97 (Sigma) plus 1 mM EDTA. After 30 min at 4 °C the insoluble material was removed by centrifugation at 12000 g and proteins were immunoprecipitated by adding specific antibodies and 30-50 µl of Protein G–Sepharose beads (Amersham Biosciences, Rainham, Essex, U.K.) to 200–2500 µl of cell lysate. After a 2-hour incubation at 4 °C under constant agitation, the beads were washed five times in lysis buffer. For reprecipitation, the proteins co-precipitated with tetraspanins were eluted in lysis buffer supplemented with 1% Triton X-100 and 0.2% SDS and identified by a second immunoprecipitation using specific antibodies. The immunoprecipitates were separated by SDS/PAGE (5–15% gel) under non-reducing conditions and transferred to a PVDF membrane (Amersham Biosciences). Biotin-labelled surface proteins were revealed using Alexa Fluor 800-labelled streptavidin (Invitrogen). The data were acquired using the Odyssey Infrared Imaging System (LI-COR Biosciences).

### Flow Cytometry

Cells were isolated as described above and incubated for the indicated time-points in the presence or absence of stimulatory factors. Cells were then collected, stained with relevant antibodies and analyzed by flow cytometry (FACSVerse BD). For intracellular flow cytometry, cells were fixed and permeabilized using the Cytofix/Cytoperm kit from BD. Cells were then stained using standard method.

### T cell proliferation assays (human and murine)

One day prior to primary cell isolation flat-bottom high binding 96-well plates (Greiner Bio-one) were coated with indicated antibody combinations in 100 μL PBS and incubated overnight at 4 °C. Plates were washed two times with PBS to eliminate unbound antibody before addition of cells. After isolation, cells were allowed to rest for 1 hour prior to staining with CellTrace Violet (Thermo Fisher Scientific) in accordance with the manufacturer’s instructions. After staining, cells were washed thoroughly with PBS before plating 1.5×105 cells to each well. Cells were then incubated for the indicated time at 37 °C, 5 % CO2, after which cells were stained for CD3, CD4 and CD8 and analyzed by flow cytometry (FACSVerse BD). FCS Express 6 software was used to analyze the proliferation of T cells based on CellTrace Violet staining. The proliferation index is defined as the average number of proliferation rounds the cells have undergone over a period of time.

### KLH immunizations and ex-vivo recall assay

Eight-week old female WT and *Cd53-/-* mice were immunized on day 0, 7 and 14 with NP-KLH (N-5060-5 Biosearch Technologies) or PBS as a control. Mice each received 100 μg NP-KLH adsorbed onto alum (Sigma) on day 0 followed by subsequent immunizations with 75 μg NP-KLH with alum per mouse on day 7 and 14. Spleens, inguinal lymph nodes and serum were harvested on day 21. Organs were processed as described in the above section, and T cells were isolated using the Pan T Cell isolation kit II according to the manufacturer’s instructions (Miltenyi Biotec). Naïve antigen presenting cells were isolated from an unimmunized (naïve) WT mouse by depleting T cells from the spleen using the CD3ε microbead kit according to the manufacturer’s instructions (Miltenyi Biotec). T cells were stained with CellTrace Violet as described in the above section and plated in 96-well flat-bottom plates at 1.5×105 T cells per well with/without 2×104 antigen presenting cells isolated from naïve WT spleen. Cells were then stimulated with medium only (control) or 100 μg NP-KLH for 96 or 120 hours, after which cells were stained for CD3, CD4 and CD8 and analyzed by flow cytometry (FACSVerse BD). FCS Express 6 was used to analyze the proliferation of CD4+ T cells based on CellTrace Violet staining and to determine the proliferation index as described above.

### IL-2 ELISA

IL-2 levels were measured using ELISA. These were performed on supernatants collected from stimulated primary T cells at the indicated time-points. For human T cells, the human IL-2 ELISA kit was used in accordance with manufacturer’s instructions (Thermo Fisher Scientific). Murine IL-2 ELISA’s were performed using the mouse IL-2 ELISA kit (Thermo Fisher Scientific).

### Micro-contact printing

Jurkat T cells were transiently transfected using the Neon transfection system according to manufacturer’s instruction, and cells were imaged 24 hours post-transfection (Thermo Fisher Scientific). Super sGFP2-CD53 was generated by subcloning hCD53 cDNA (NM_000560, Thermo Scientific) into psGFP-C118. The construct encoding human CD37 was previously described and subcloned into psGFP2-C1 vector_19_. mCherry-PKCθ was generated by subcloning of hPKCθ (GenScript) into mCherry-N120. Poly(dimethylsiloxane) stamps containing a regular pattern of circular spots with a diameter of 5 μm were prepared as described previously_21, 22_. Stamps were incubated for 1 hour with phosphate-buffered saline (PBS) containing anti-CD3 and anti-CD28 antibodies or isotype control antibodies (100 μg/ml) mixed with donkey anti-rabbit IgG (H&L)– Alexa Fluor 647 (10 μg/ml, Invitrogen) to visualize the spots. Stamps were washed with demineralized water and dried under a nitrogen stream. The stamp was applied to a cleaned glass coverslip for 20 s and then removed. Transfected cells were seeded on the stamped area and incubated at 37°C for 5 min. Paraformaldehyde (PFA) was added to the cells to a final concentration of 2 % PFA, or the coverslips were washed with PBS and fixed with 2 % PFA in PBS for 20 min at room temperature. Samples were washed with PBS and demineralized water and embedded in Mowiol (Sigma). Cells were imaged on an epifluorescence Leica DMI6000 microscope with a 63× oil 1.4 NA objective, a metal halide EL6000 lamp for excitation, a DFC365 FX CCD camera (Leica), and GFP, DsRed, and Y5 filter sets (for GFP, RFP, and Alexa Fluor 647, respectively; all from Leica). Focus was kept stable with the adaptive focus control from Leica.

### Transfection and Fluorescence Lifetime Imaging Microscopy (FLIM)

Jurkat T cells were transiently transfected using the Neon transfection system according to manufacturer’s instruction, and cells were imaged 24 hours post-transfection (Thermo Fisher Scientific). mCitrine-CD53 was generated by subcloning hCD53 cDNA (NM_000560, Thermo Scientific) into pmCitrine-C123. mCherry-PKCθ was generated by subcloning of hPKCθ (GenScript) into mCherry-N120. Phorbol 12-myristate 13-acetate (PMA) was used to induce the translocation of PKCθ to the membrane. Live-cell microscopic analysis for FLIM in Jurkat T cells was performed on a SP8 confocal microscope equipped with a 60× water 1.2 NA objective (Leica) using appropriate laser lines and settings. FLIM images were recorded using the same setup with an excitation light at 495 nm provided by a pulsed white-light laser (Leica). Fluorescence (from 500 to 535 nm) was collected with an internal photomultiplier tube and processed by a PicoHarp 300 time-correlated single-photon counting system (PicoQuant). At least 50,000 photons were recorded for each individual cell.

To obtain the fluorescence lifetimes, photon histograms containing all photons pooled for each individual cell were reconstructed from the photon traces with a custom programmed algorithm (C#.NET), and these histograms were fitted with single-exponential decay curves (OriginLab) to obtain donor fluorescence lifetimes. A biexponential fit was used to obtain the percentages of bound donor. The typical lifetime of the donor was a fixed fitting parameter in this analysis.

### Generation of human CD53-deficient CEM T cells by CRISPR/Cas9-mediated genome editing

One guide RNA pair targeting sequences TGATAGAGCCCATGCAGCCCAGG and GTGAGTCCTTACAGCAGATGTGG within the CD53 gene were designed using the Massachusetts Institute Technology CRISPR design tool_24_ and subcloned into the px335 Cas9 vector as described before_25_. CEM T cells were seeded on day 0 at an optimal density of 3 x105 cells/ml. On day 1 cells were transfected with 1 μg of each of the two guide RNA plasmids and 0.5 ug of pGFP2-C1 vector with a Neon Transfection System Kit and Neon Transfection System (Thermo Fisher Scientific) according to manufacturer’s guidelines as described for this cell line. On day 3, CEM cells were selected by fluorescence-activated cell sorting based on GFP expression using FACS Aria III (BD Bioscience). Eight days later, CD53-negative CEM cells were selected by fluorescence-activated cell sorting using an AF488-conjugated antibody against CD53 (MEM53, Novus Biologicals). CD53 deficiency was validated by flow cytometry and Western blotting.

### FRAP of WT and CD53-/-CEM T cells

WT and *CD53-/-* CEM T cells were transfected using the Neon transfection system (Thermo Fisher Scientific) with 2 µg of plasmid encoding for human CD45RO-GFP. CD45RO construct was made to order by Genscript and re-cloned in house into a psGFP2-N1 vector. 24 hours post-transfection cells were seeded in Willco dishes and FRAP was performed using a Leica TCS SP8 SMD microscope equipped with a 60× water 1.2 NA objective (Leica) and an argon-ion laser set to bleach with 40 % power at the 488 nm wavelength. We measured the fluorescence intensity in the bleach zone as well as the whole cell and background in order to correct for photobleaching and background signal. Immobile and mobile fractions were calculated manually and confirmed using the easyFRAP web tool_26_. Fitting of the average curve was performed with OriginPro 8 (Originlab).

### CD45 Internalization Assay

*CD53-/-* and WT CEM T cells were transfected using the Neon transfection system (Thermo Fisher Scientific) with 2ug of plasmid encoding for human CD45RO-GFP. 24 hours post-transfection, cells were labeled with an anti-CD45RO antibody and incubated at 37°C (5% CO2) for 0, 3, 6 and 24 hours post-labeling. Upon harvesting, cells were fixed in 4% PFA, and stained with a secondary Alexa-647 antibody to quantify the amount of CD45RO antibody remaining on the surface. Cells were then measured using flow cytometry (FACSVerse BD), and analysis was performed using the FlowJo software package.

### Statistical analysis

All statistical comparisons were made with GraphPad Prism 5 software, and data are expressed as means ± SEM, SD, or 95% CI as indicated in the figure legends. Differences between means were analyzed with Student’s t tests or two-way ANOVA as indicated in the legends. Statistical significance was set at P < 0.05.

