## Supplementary Figures for "Dynamic regulation of CD45 by tetraspanin CD53"

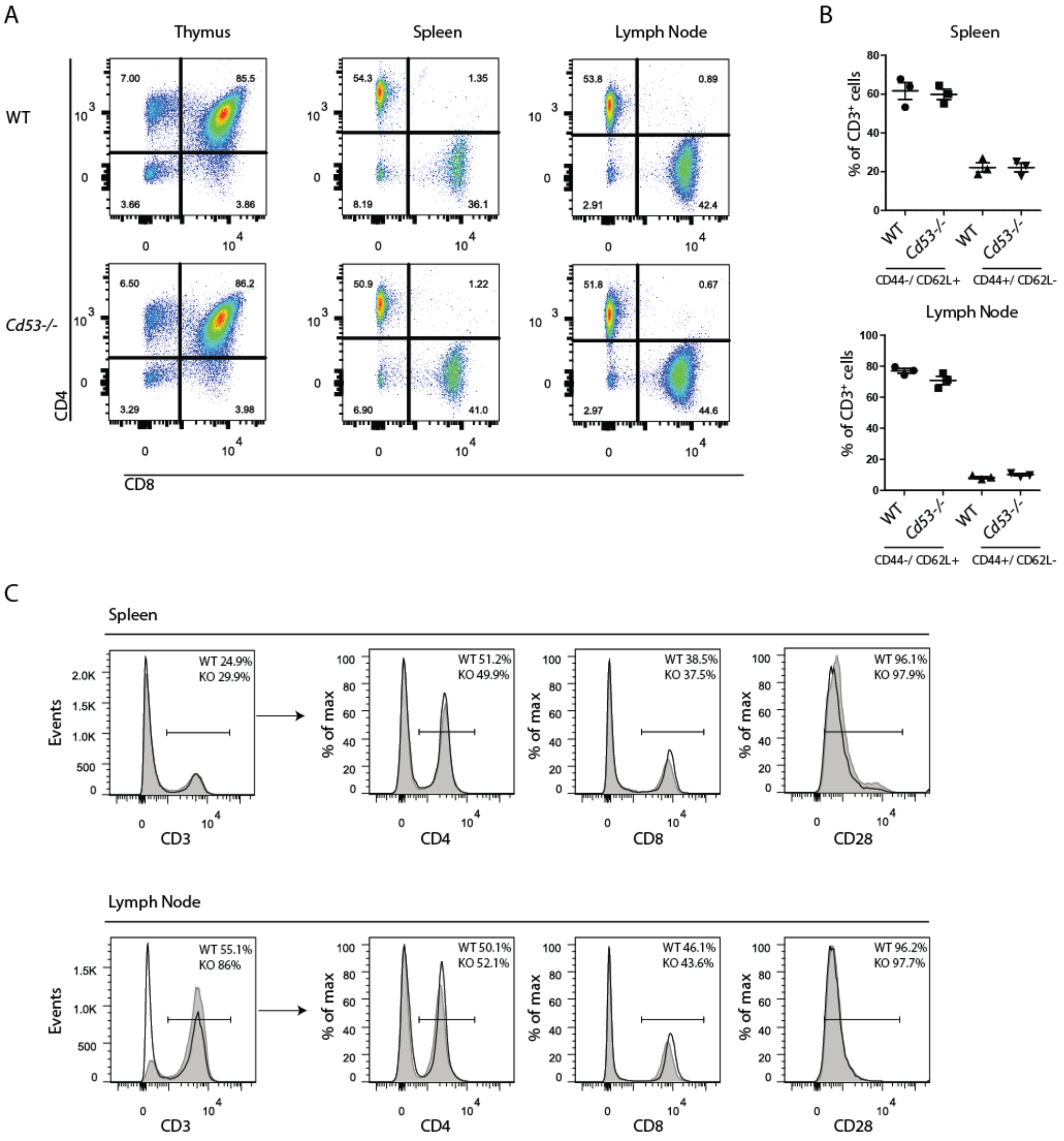


**Figure S1. T cell development is normal in *Cd53-/-* mice.**
**(A)** two-parameter dot plots showing the CD4 and CD8 expression of cells from the thymus (left), spleen (middle) and lymph nodes (right) of six week old WT and *Cd53-/-* mice as measured by flow cytometry. For thymus all cells isolated from the organ were included. For spleen and lymph node, cells were first gated based on CD3 expression. Data was gathered from three mice per genotype from one independent experiment. Representative data from one WT and one Cd53-/- mouse is shown. **(B)** Percentage of memory (CD44+/CD62L-) and naive (CD44-/CD62L+) T cells within the CD3+ population in spleen and lymph nodes for both WT and *Cd53-/-* mice. Data was gathered from three mice per genotype from one independent experiment. Statistical significance was assessed by unpaired t test. **(C)** Expression of CD3, CD4, CD8 and CD28 as measured by flow cytometry for three mice per genotype from one independent experiment. Representative data from one WT and one *Cd53-/-* mouse is shown. Percentage gates are set based on unstained samples (not shown).

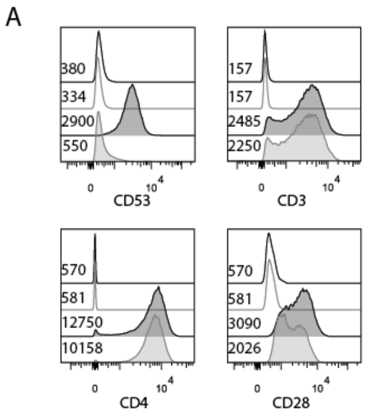


**Figure S1. Expression of CD53, CD4, CD8 and CD28 on WT and *CD53-/-* CEM T cells.**
**(A)** Expression of CD53, CD3, CD4 and CD28 on WT and *CD53-/-* CEM T cells. Unfilled black and gray lines represent unstained WT and *CD53-/-* CEM T cells, respectively. Filled black and gray graphs represent stained WT and *CD53-/-* CEM T cells, respectively. Corresponding gMFI values are indicated at the left side of each graph.

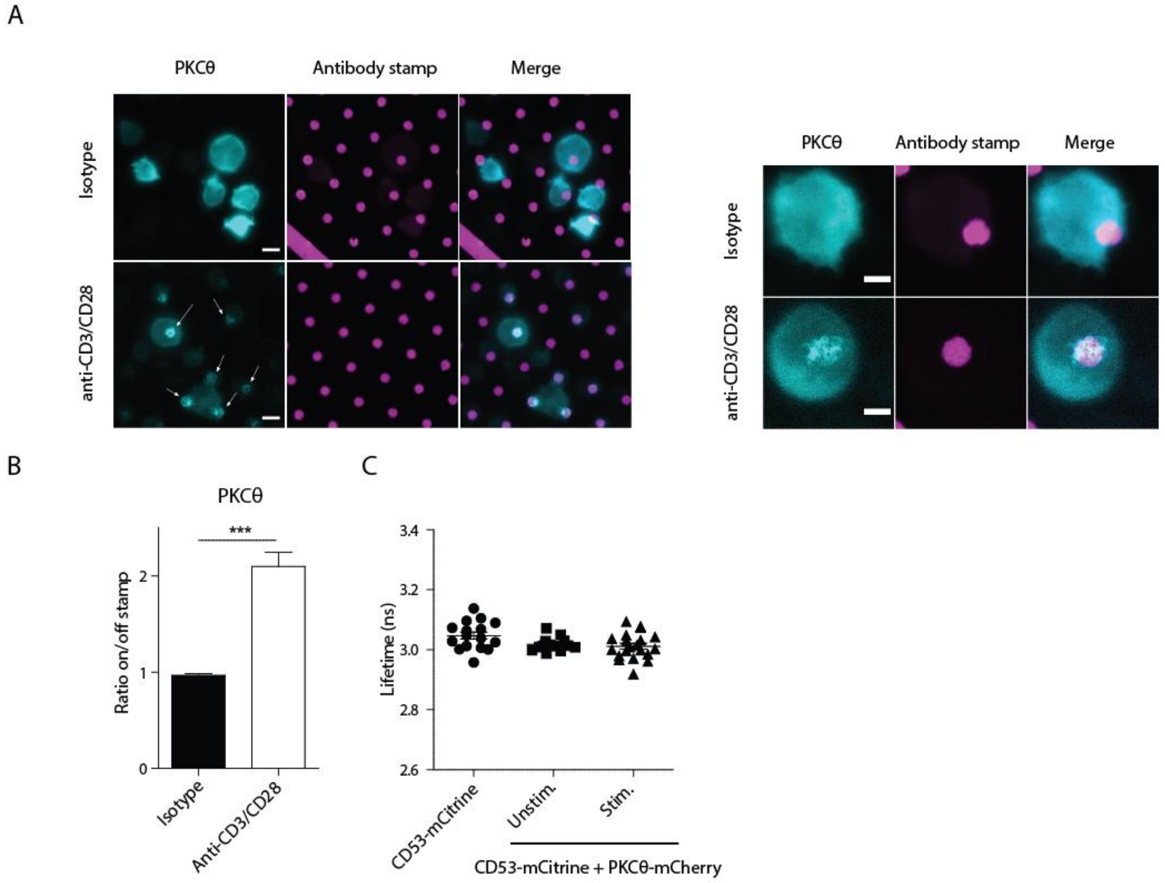


**Figure S3. PKCθ is recruited to TCR signaling sites, where it does not interact with CD53.**
**(A)** Jurkat T cells expressing PKCθ -mCherry were seeded on isotype control or anti-CD3/CD28 prints and PKCθ recruitment was analyzed after 8 minutes by epifluorescence microscopy. Left: overview of multiple cells. Right: larger magnification of one representative cell. **(B)** Ratio between the average PKCθ intensity on the antibody print and that outside of the print. Data represents analysis of >30 cells from three independent experiments. Statistical significance was assessed by one-sample t test. **(C)** Jurkat T cells experssing CD53-mCitrine alone, or coexpressing CD53-mCitrine and PKC-mCherry were subjected to FLIM analysis. Graph shows lifetimes of CD53-mCitrine obtained by fitting fluorescent decay curves with mono-exponential decay functions. Data are means ± SEM of at least 14 cells per condition from one experiment. Statistical significance was assessed by unpaired t test. Data are means ± SEM (* p<0.05, ** p>0.01, *** p> 0.001). Scale bars represent either 10 µm (left panels) or 5 µm (right panels) as indicated in the bottom right corner of the leftmost images. Arrows indicate TCR signaling sites showing PKCθ recruitment.
